# Silencing of hippocampal synaptic transmission impairs spatial reward search on a head-fixed tactile treadmill task

**DOI:** 10.1101/2021.09.03.458092

**Authors:** Jake T. Jordan, J. Tiago Gonçalves

## Abstract

Head-fixed linear treadmill tasks have been used to study hippocampal physiology in mice. Although some hippocampal neurons establish place fields along linear treadmills, it is not clear if the hippocampus is required for spatial memory on this task. Using a Designer Receptors Exclusively Activated by Designer Drugs (DREADDs) approach, we found that silencing hippocampal output on rewarded treadmill tasks impaired search for rewards signaled by spatial cues but did not impair search for rewards signaled by local cues, recapitulating findings from other behavior tasks. These findings serve to contextualize data on hippocampal physiology from mice performing this task.

## Introduction

Spatial coding in the rodent hippocampus has been studied extensively in a large variety of spatial contexts. In many experiments, rewards are used to motivate the animal to move around an environment. Both rewards and attentional demands have been shown to shape place cell activity and stability (Kentros et al., 2004; Fenton et al., 2010; Gauthier & Tank, 2018), however, in many cases the animals may not need to use spatial cues to find the rewards and it is unclear how hippocampal representations may be affected by whether a task requires hippocampal activity.

Linear environments enable the recording of mouse hippocampal activity through many traversals of each location within a session and some hippocampal neurons will establish place fields along the linear environment (Harvey et al., 2009; Dombeck et al., 2010; Royer et al., 2012; Ziv et al., 2013; Bittner et al., 2015 and 2017; Sheffield et al., 2015; Danielson et al., 2016; Grienberger et al., 2017; Sato et al., 2017 and 2020; Sheffield & Dombeck, 2017; Zaremba et al., 2017; Hainmueller & Bartos, 2018; Hayashi, 2018; Gonzalez et al., 2019; Lee et al., 2020; Robinson et al., 2021; Mizuta et al., 2021). A variety of linear environment tasks have been used, most of which involve administering a water reward to a water-deprived mouse at a certain location within the environment. However, these tasks differ in how rewards are administered and whether spatial cues need to be attended to identify rewarded locations. As rewards shape spatial representations (Gauthier & Tank, 2018), differences in reward contingencies may affect whether the hippocampus is required for performance of a task and may even affect the long-term dynamics of hippocampal representations. Further, attentional demands affect both place cell stability (Kentros et al., 2004) and the spatial distribution of place cell firing (Fenton et al., 2010). Thus, in order to best interpret physiological findings from a particular behavioral task, it should be determined whether the hippocampus is required for task performance.

Head-fixed spatial memory tasks involve the administration of “hidden” water rewards at fixed locations. In a head-fixed virtual reality (VR) linear treadmill task using visual cues, mutations that affect hippocampal place cell dynamics and inhibition of the dorsal hippocampus impaired search for the location of a water reward (Sato et al., 2017 and 2020) and stimulation of place cells is sufficient to manipulate spatial reward search (Robinson et al., 2021) suggesting that hippocampal activity recorded during performance of this task reflects the neural dynamics that are in some way involved in memory retrieval. In a similar head-fixed linear treadmill task that makes use of a tactile treadmill belt instead of a VR simulation, genetic mutations (Zaremba et al., 2017) or manipulations of hippocampal interneuron activity (Turi et al., 2019) that affect hippocampal place cell dynamics have been shown not to impair spatial memory for rewards at a single location as measured by the spatial distribution of lick behavior (Zaremba et al., 2017; Turi et al., 2019). However, when required to learn a new location, reward search was impaired, suggesting that natural place cell dynamics are not necessary for initial learning but are required for learning multiple environments (Zaremba et al., 2017; Turi et al., 2019). It is not known if activity in hippocampal pyramidal neurons is required for memory of a single reward location on the tactile linear treadmill task. Determining whether this is the case will help to interpret data on hippocampal physiology during performance of this task.

Here, we used two versions of the tactile linear treadmill task in which water-restricted mice received water rewards at a particular location along the treadmill belt (Figure 1). In one version, this location was signaled by salient local cues. In other behavioral paradigms, such as the Morris Water Maze (MWM), memory for locally-cued goal locations is insensitive to hippocampal damage (Morris et al., 1982). In the second version of the task, reward location was signaled by non-local spatial cues. This type of goal location memory has been shown to be impaired by hippocampal damage in the MWM (Morris et al., 1982). We then tested how reward location memory for both versions of this task was affected by silencing of dorsal hippocampal output.

**Figure 1.**
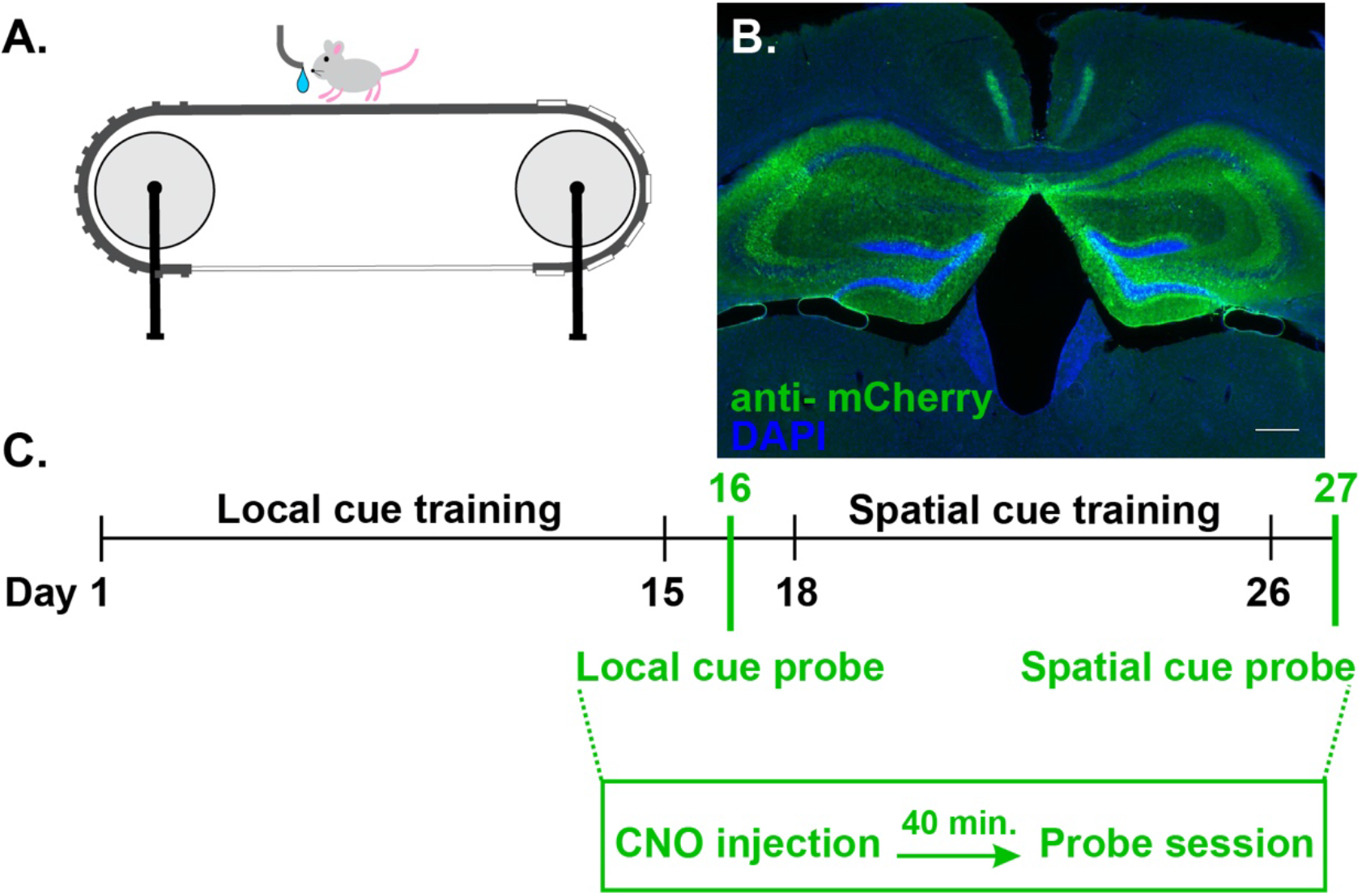
Experimental design. (a) Illustration of treadmill water reward task. Mice remain stationary via head-fixation as they pull the treadmill belt. At a specific location along the belt, a water reward is administered. A sensor attached to the reward spout acquires lick data. (b) A coronal section of the bilateral dorsal hippocampus injected with a virus to drive expression of hM4D_Gi_ fused to mCherry. Expression of the fusion construct is shown here as an immunostaining using an antibody against mCherry. Control mice expressed mCherry only. Scalebar = 250 μm (c) Timeline of experiments.

## Results

We injected the dorsal hippocampus of experimental mice with a virus to drive expression of hM4D_Gi_ conjugated to mCherry and control mice with a virus to drive expression of mCherry alone (Figure 1b). Activation of hM4D_Gi_ via systemic injection of CNO silences synaptic transmission (Stachniak et al., 2014) and doing so in the hippocampus impairs hippocampus-dependent memory (Varela et al., 2016). Mice were water restricted using 2%citric acid water (Urai et al., 2020) and were then trained to run on the treadmill for water rewards (Figure 1c). To assess learning and memory of the reward location, we used a lick sensor to quantify the fraction of licks occurring near the reward location (Danielson et al., 2016; Zaremba et al., 2017; Turi et al., 2019).

We first sought to test whether hippocampal silencing would impair reward location memory signaled by a local cue (Figure 2a). Based on studies of rodent hippocampal function using other behavioral paradigms, such as the Morris Water Maze, we predicted that hippocampal silencing would not affect this type of behavior. Both groups of mice earned a similar number of rewards during cued reward learning (day: p < 0.0001; virus: p = 0.819; interaction: p = 0.510, Figure 2b). Mice expressing hM4D_Gi_ learned to lick selectively near the reward location faster than control mice (day: p = 0.008; virus: p = 0.041, interaction: p = 0.203), however, this appeared to be driven by early sessions when some mice were not yet completing a full lap and therefore the fraction of licks near the reward zone could not be computed and sample sizes were incomplete. Excluding the first week, both groups of mice showed a similar degree of reward zone lick selectivity (day: p = 0.001; virus: p = 0.381; interaction: p = 0.588, Figure 2c). To test reward location memory, a probe session was run where rewards were not administered and the hippocampus was inhibited via injection of CNO 40 minutes prior to the beginning of the session (Figure 2d). There was no effect of virus on the number of laps run in the probe session (p = 0.250, Figure 2e). Despite withdrawal of reward, mice not only licked the reward spout, but licked selectively in the reward location. Lick rates did not differ between groups (p = 0.569). Further, the fraction of licks in the rewarded zone were much higher than chance (p < 0.0001 for both groups) but did not differ between groups (p = 0.293, Figure 2f), despite the absence of reward. In line with our prediction, these data suggest that hippocampal silencing does not affect memory of reward locations signaled by a local cue.

**Figure 2.**
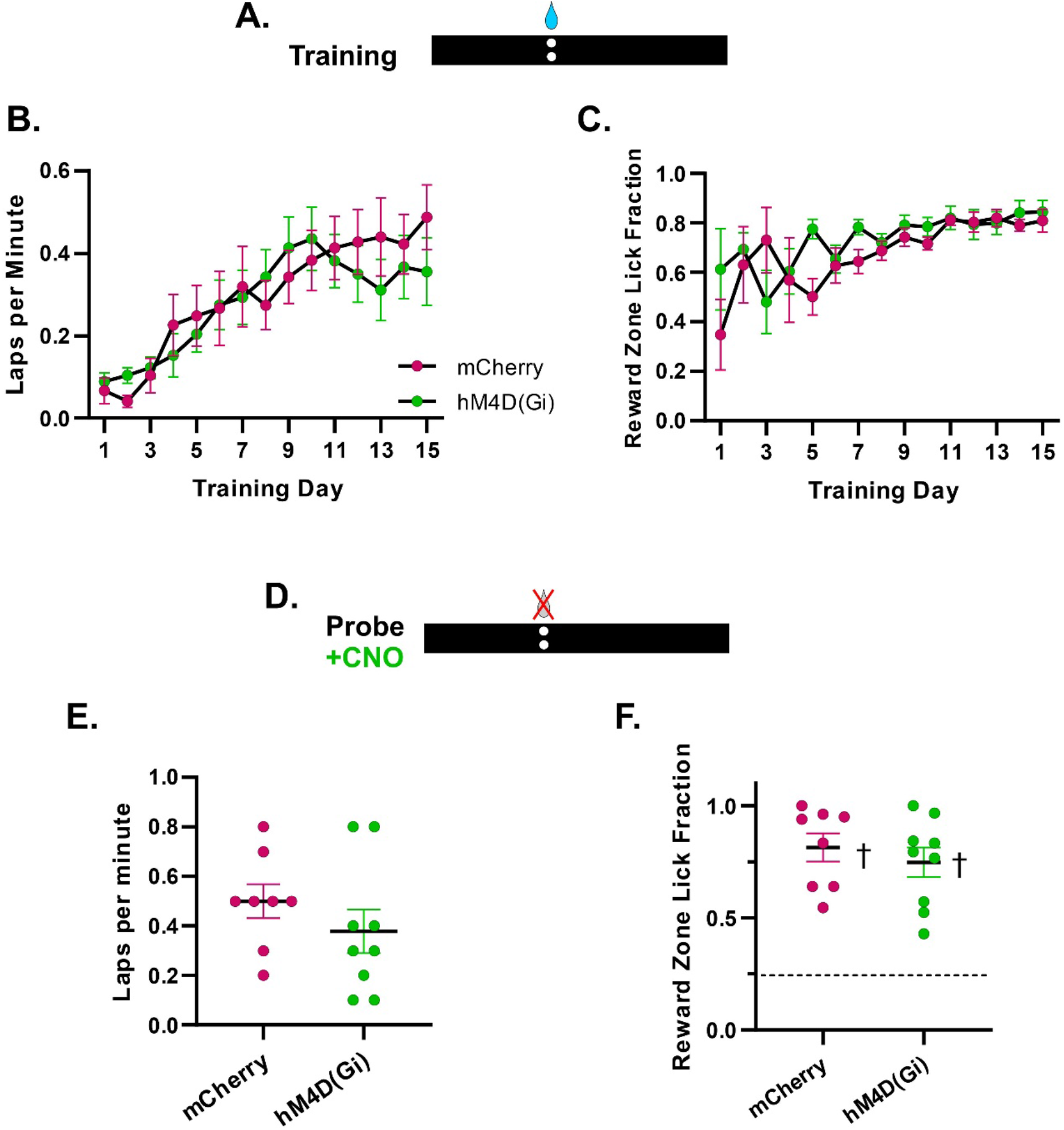
Dorsal hippocampal silencing did not impair locally-cued search for reward memories. (a) During training, reward location was signaled by a local salient cue at the reward site. (b) Number of laps completed per minute during treadmill training. (c) The fraction of all licks of the reward spout that occurred within the rewarded quadrant of the treadmill belt during training. (d) Mice were injected with CNO and run on a probe trial 40 minutes later in which rewards were omitted. (e) Number of laps completed per minute during the probe session. (f) The fraction of all licks of the reward spout that occurred within the rewarded quadrant of the treadmill belt during the probe session. † indicates group is significantly different from chance performance, p < 0.0001, one-sample t-test with Bonferroni’s correction.

We sought to test whether hippocampal silencing would impair reward location memory signaled by non-local spatial cues (Figure 3a). Both groups of mice earned a similar number of rewards during spatial learning (day: p = 0.146; virus: p = 0.445; interaction: p = 0.036, Figure 3b). Despite a day by virus interaction, reward rates did not differ between groups on any day (p ≥ 0.59). Both groups of mice licked selectively near the rewarded location but did not differ from each other (day: p = 0.006; virus: p = 0.797; interaction: p = 0.162, Figure 3c). A probe session was run where rewards were not administered and the hippocampus was silenced via injection of CNO 40 minutes prior to the beginning of the session. There was no effect of virus on the number of laps run (p = 0.122) or lick rates (p = 0.105, Figure 3e) during the probe session. However, the fraction of licks in the rewarded zone was lower in the hM4D_Gi_ group compared to controls (p = 0.034, Figure 3f). Further, while control mice licked selectively near the rewarded location (p = 0.015), mice expressing hM4D_Gi_ did not (p = 0.076). In line with our prediction, these data suggest that hippocampal silencing impairs search at a previously rewarded location using non-local spatial cues.

**Figure 3.**
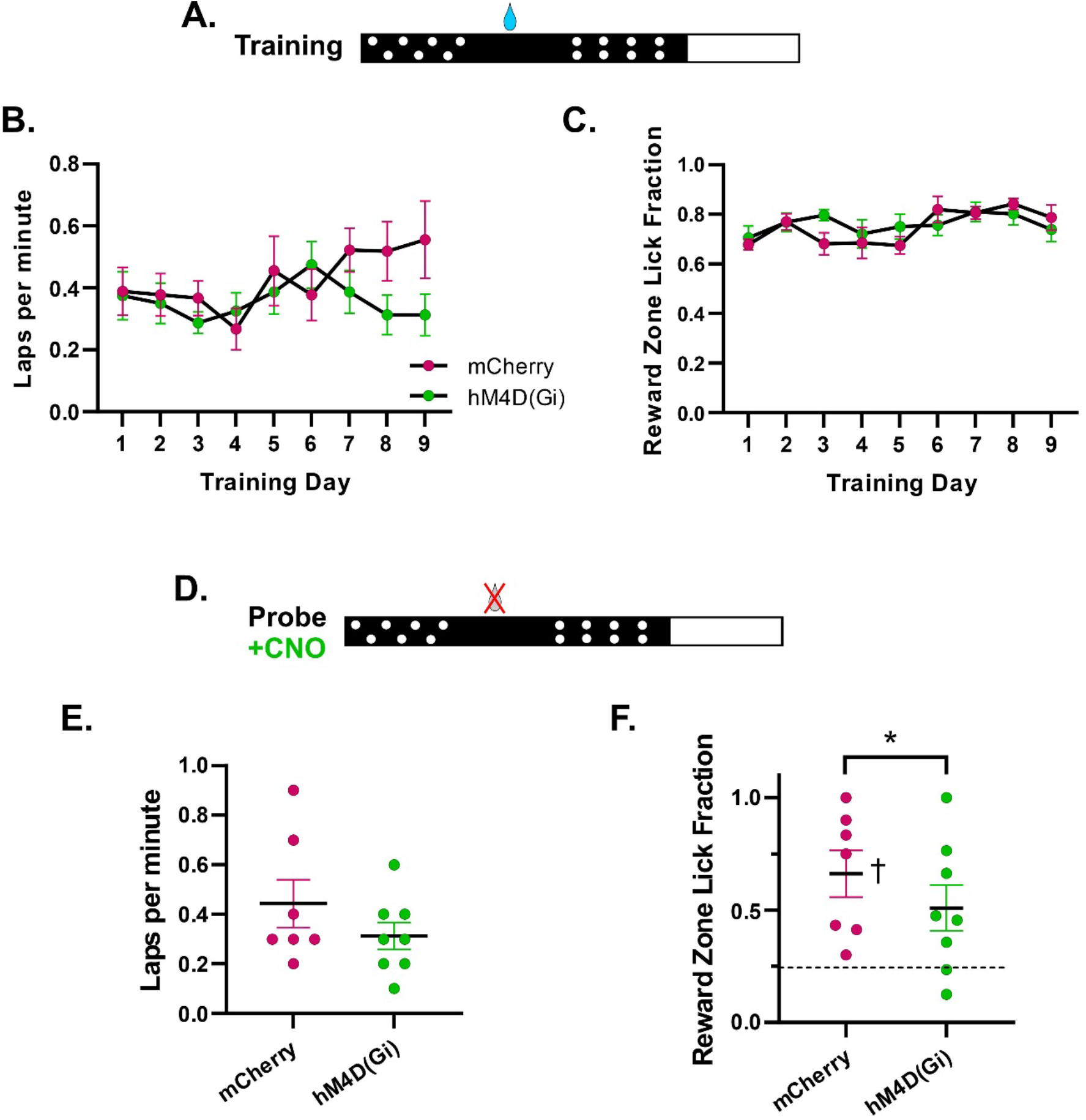
Dorsal hippocampal silencing impairs spatial search for reward memories. (a) During training, reward location was signaled by spatial cues along the treadmill belt. (b) Number of laps completed per minute during treadmill training. (c) The fraction of all licks of the reward spout that occurred within the rewarded quadrant of the treadmill belt during training. (d) Mice were injected with CNO and run on a probe trial 40 minutes later in which rewards were omitted. (e) Number of laps completed per minute during the probe session. (f) The fraction of all licks of the reward spout that occurred within the rewarded quadrant of the treadmill belt during the probe session. * indicates hM4D_Gi_ mice differed from controls, p < 0.05; † indicates group is significantly different from chance performance, p < 0.0001.

## Discussion

Head-fixed tactile linear treadmill tasks have been used to identify neural phenomena associated with spatial learning and memory (Bittner et al., 2015; Danielson et al., 2016; Bittner et al., 2017; Grienberger et al., 2017; Zaremba et al., 2017; Turi et al., 2019). However, it was previously unknown if dorsal hippocampal activity was required for retrieval of rewarded locations. Here, we have found that silencing of hippocampal output via hM4D_Gi_ activation impaired search for a reward location when using spatial cues. It is unlikely this is due to a change in motivation as experimental and control groups ran a similar number of laps and licked the reward spout at a similar rate. Further, it is unlikely that this is due to a more general memory impairment as there was no impairment of search for reward locations signaled by local cues, in line with studies using other behavioral paradigms, such as the MWM (Morris et al., 1982) or the T-Maze (Shipton et al., 2014). Thus, spatial memories for reward locations on a linear treadmill depend on output of the dorsal hippocampus.

Chromosomal deletions and manipulations of hippocampal interneuron activity that alter hippocampal spatial representations have been shown to impair learning of multiple reward locations while sparing learning of a single reward location (Zaremba et al., 2017; Turi et al., 2019). In the context of our current study, it appears that memory for a single rewarded location on a tactile treadmill requires activity in dorsal hippocampal pyramidal cells, even if that activity is altered. It is not clear at this time how brain regions which interact with the hippocampus work to compensate for altered hippocampal activity and why this fails when learning multiple environments.

Recording of neuronal activity via calcium imaging has enabled long-term recording of hippocampal neurons across many days, weeks, and even months (Dombeck et al., 2010; Ziv et al., 2013; Rubin et al., 2015; Hainmueller & Bartos, 2018; Hayashi, 2018; Gonzalez et al., 2019; Lee et al., 2020; Robinson et al., 2021; Sato et al., 2020; Sheintuch et al., 2020; Mizuta et al., 2021). These studies have found that the population of neurons encoding a particular space evolve gradually over days and weeks (Ziv et al., 2013; Rubin et al., 2015; Danielson et al., 2016; Zaremba et al., 2017; Hainmueller & Bartos, 2018; Hayashi, 2018; Gonzalez et al., 2019; Lee et al., 2020; Sato et al., 2020; Sheintuch et al., 2020; Mizuta et al., 2021). This has led to some interesting theories regarding what the functions are of the turnover of the active population (Hayashi, 2018; Mau et al., 2020) and whether stable versus non-stable place cells offer unique contributions to learning and memory (Lee et al., 2020; Mau et al., 2020). One limitation in generalizing findings across studies is differences in behavioral tasks. For instance, “linear track” tasks, in which mice run from end of a track to another and receive a water reward at the end of the track, can be completed despite hippocampal damage (Gonzalez et al., 2019) and does not require attention to spatial cues. Interestingly, the hippocampus has been shown to spontaneously remap between recording sessions despite no changes to the environment or spatial cues (Sheintuch et al., 2020), suggesting the possibility that the hippocampus contains multiple maps of single environments. However, this remapping could be partially accounted for by 180° rotations of hippocampal spatial representations relative to the environment (Gonzalez et al., 2019; Sheintuch et al., 2020). It is not clear whether increased task demands would also result in multiple representations of a fixed environment. On the other hand, dorsal hippocampal activity is required when searching for hidden water rewards in a VR environment (Sato et al., 2017) and as we have shown when using tactile spatial cues. Studies using these tasks have identified similar neural phenomena as studies using the linear track task, e.g. gradual turnover of the hippocampal population code for a given environment (Ziv et al., 2013; Lee et al., 2020). However, it remains possible that recording hippocampal activity during performance of one type of task may result in different neuronal dynamics than would performance of another task.

## Acknowledgments

The authors wish to thank Sunaina Martin and Cesar Diaz for assistance with histology as well as Agustina Frechou and Kelsey McDermott for comments on the manuscript. J.T.J was funded by an NIH T32 training grant (T32HD098067) and would like to thank the Intellectual and Developmental Disabilities Research Center for its support (1 P50 HD105352-01; support for the Rose F. Kennedy IDDRC). J.T.G. was funded by a Whitehall Foundation Research Grant (2019-05-71).

## Methods

### Animals

Male and female C57BL6/J mice (Jackson Labs and bred in-house) were kept on a 12h light/dark cycle and were allowed standard chow *ad libitum.* Water was available *ad libitum* prior to the beginning of the experiment. Mice were housed in groups of 3-5 and littermates were divided between experimental groups. All procedures were done with approval of the Institutional Animal Care and Use Committee (protocol #: 00001197).

### Surgery

Mice, aged 8-11 weeks at time of surgery, were deeply anesthetized with 5% isoflurane and were then kept under anesthesia with an oxygen flow (0.5 L per minute) and 2% isoflurane. Bilateral hippocampus (AP: 1.85 mm, ML: ±1.7 mm, DV: 1.45mm) was stereotactically injected with either AAV8-CaMKIIα-hM4D_Gi_-mCherry (experimental mice: n = 9; 5 males, 4 females) or AAV5-CaMKIIα-mCherry (controls: n = 9; 2 males, 7 females) at a titer of 1 × 10^12^ GC/ml. A small (<1 mm) tungsten wire soldered to a gold pin (used to ground the mice for lick counting) was implanted above the cerebellum and a metal headbar was affixed to mouse’s skull using dental cement. Mice were given Carprofen (5mg/Kg) and were allowed one week of recovery before water restriction began.

### Pharmacology

The hM4D_Gi_ agonist clozapine-N-oxide (CNO) was dissolved at a concentration of 1 mg/mL in saline (0.9% NaCl) and was injected intraperitoneally at a dose of 5 mg/kg (as done by Varela et al., 2016). CNO was injected 40 minutes prior to behavior sessions.

### Water restriction with 2% citric acid water

The mass of each mouse was recorded both prior to citric acid water treatment, in the immediate four days after beginning citric acid water treatment, and at least once per week afterwards. One week prior to behavioral training, homecage water was replaced with water containing 2% citric acid (Urai et al., 2020). A threshold of 15% weight loss was used to determine if mice were healthy enough to remain on water restriction. No mouse fell below this weight.

### Linear treadmill apparatus

The 180 cm linear treadmill apparatus was similar to those previously used (Royer et al., 2012). Tasks were controlled with custom MATLAB software and data from the treadmill were acquired using National Instruments data acquisition hardware. The treadmill consisted of a velvet-textured belt wound around two wheels. An optical rotary encoder was attached to the axel of one wheel to measure forward and backward movement of the belt, enabling the estimation of the position of the mouse along the belt. Four radio frequency identification (RFID) tags were attached to the belt at regular intervals (every 45 cm) to correct the accumulating error of position estimation by the rotary encoder data. The RFID scanner was affixed to the mouse platform. For spatial tasks, RFID tags were attached to the belt at transition zones between textures. Additional RFID tags were used to trigger reward delivery (1 per lap; 10 μL of water; water contained 10% artificial sweetener) through a microinjection pump controlled by TTL pulses from the National Instruments hardware. Rewards could only be delivered at a single location when a reward RFID tag was scanned in sequence with the other RFID tag in front of it on the belt. Reward seeking was measured with a lick counter (Janelia Farm, VA, USA). This prevents reward delivery during backward walking and reinforces forward movement during the initial training phase.

### Habituation and behavioral training

Mice were habituated to the experimenter and environment and were trained to run for water rewards as described previously (Jordan et al., 2021). In the initial phase of training during which mice learned to associate reward location with a single local cue, the treadmill belt had spatially uniform stripes made with a hot glue gun (so that the mouse could grip the belt) and a single pair of sandpaper disks (100 grit, 2.5 cm diameter) 10 cm in front of the site of reward delivery. For spatial memory tasks, belts with four tactile zones were used. Rewards were administered near the middle of one of these zones, approximately 19 cm away from the transition between the reward zone and the previous zone and from any tactile cues. We first trained mice to learn the location of a hidden water reward. After 9 days of training, mice were given a single probe trial in which rewards were omitted and bilateral dorsal hippocampus was silenced via injection of CNO. Spatial memory was measured by the fraction of all licks occurring in the reward zone (Danielson et al., 2016; Zaremba et al., 2017; Turi et al., 2019). Probe trials were run by an experimenter blind to treatment group.

### Statistical analysis

Power analysis was done following a pilot cohort. A power analysis was conducted on the first cohort of mice (5 experimental, 5 control) which revealed that sample sizes of 8 mice per group would yield 80% power. Thus, we added a second cohort of mice which yield a final sample size of 9 mice per group. There were no differences across sexes and data from males and females were combined (Shansky, 2019). One control mouse was not run on the cued probe session due to distress. One mouse was removed from the spatial memory experiment as the grounding pin required for lick detection fell out. Two control mice were removed from the spatial probe analysis: one did not lick at all during the probe session and the other did not complete any laps, making it impossible to calculate a reward zone lick fraction. Two-way mixed model ANOVAs were used to compare performance of the groups across acquisition. Post hoc analyses were done using Sidak’s multiple comparisons test. To determine whether behavior differed between experimental and control mice during the probe session, linear mixed effects models were used which accounted for differences between the first and second cohorts (see above). One-sample t-tests with Bonferroni’s correction for multiple comparisons were run to determine differences between group means and chance performance.

### Immunohistochemistry

Animals were perfused with ice cold 0.1M phosphate buffer saline followed by 4% paraformaldehyde (PFA) and were then post-fixed in 4% PFA for 24 hours. Brains were then cryoprotected in 30% sucrose and the hippocampus was sectioned at 40 μm thickness on a freezing microtome. Sections were rinsed three times in 0.1M PBS (pH = 7.4) and incubated in a blocking solution (10% goat serum, 0.3% Triton-X in 0.1M PBS) for one hour. The sectioned tissue was then incubated in a rabbit anti-mCherry primary antibody in block (1:1000; EnCor Biotechnology; Lot #: 205-120820) for 48 hours, followed by three rinses in 0.1M PBS. The sections were then incubated in Alexa 488 goat anti-rabbit secondary in 0.1M PBS (1:200; Invitrogen; Lot #: 2286890) for 2 hours, rinsed three times in 0.1M PBS, counterstained with 300nM DAPI in 0.1M PBS, rinsed again in 0.1M PBS, and then mounted and cover-slipped with Fluoromount-G (SouthernBiotech; Lot #: H0120-V110). The tissue was then imaged on a Zeiss Axio Imager.A2 fluorescence microscope. Animals in which there was no expression in the dorsal hippocampus were removed from analysis.

